# Membrane Protrusion Formation Mediated by Rho/ROCK Signalling and Modulation of Chloride Flux

**DOI:** 10.1101/600379

**Authors:** Akiko Hori, Kenji Nishide, Yuki Yasukuni, Kei Haga, Wataru Kakuta, Yasuyuki Ishikawa, Matthew J Hayes, Shin-ichi Ohnuma, Hiroshi Kiyonari, Kazuhiro Kimura, Toru Kondo, Noriaki Sasai

**Affiliations:** Developmental Biomedical Science, Division of Biological Sciences, Nara Institute of Science and Technology, 8916-5, Takayama-cho, Ikoma 630-0192, Japan; Laboratory for Cell Lineage Modulation, Centre for Developmental Biology, 2-2-3, Minatojima-Minamimachi, Chuo-ku, Kobe 650-0047, Japan; Department of Systems Life Engineering, Maebashi Institute of Technology, 460-1, Kamisadori-cho, Maebashi, Gunma, 371-0816, Japan; Ocular Biology and Therapeutics Unit (ORBIT), Faculty of Brain Sciences, UCL Institute of Ophthalmology, University College London, 11-43 Bath Street, London EC1V 9EL, United Kingdom; Laboratory for Animal Resources and Genetic Engineering, RIKEN Center for Biosystems Dynamics Research, 2-2-3, Minatojima-Minamimachi, Chuo-ku, Kobe 650-0047, Japan; Department of Ophthalmology, Yamaguchi University School of Medicine, 1-1-1 Minamikogushi, Ube 755-0046, Japan; Division of Stem Cell Biology, Institute for Genetic Medicine, Hokkaido University, Kita-15, Nishi-7, Kita-Ku, Sapporo 060-0815, Japan

## Abstract

Membrane protrusion is an important structural property associated with various cellular functions. The pentaspan membrane protein Prominin-1 (Prom1/CD133) is known to be localised to the protrusions and plays a pivotal role in migration and the determination of cellular morphology; however, the underlying mechanisms have been elusive. Here, we demonstrate that Prom1 is sufficient to trigger membrane protrusion formation. Overexpression of Prom1 in the RPE-1 cells triggers multiple long cholesterol-enriched protrusions, independently from actin and tubulin polymerisation. For this protrusion formation, the five amino acid stretch located at the carboxyl cytosolic region is essential. Moreover, the small GTPase Rho and its effector kinase ROCK are essential for this protrusion formation, and the intersection point of active Rho and Prom1 is where the protrusion formation initiates. Importantly, Prom1 causes the chloride ion efflux induced by calcium ion uptake, and protrusion formation is closely associated with the chloride efflux activity. Altogether, this study has elucidated that Prom1 plays critical roles for the membrane morphology and chloride ion flux.

## INTRODUCTION

Each cell has a unique shape corresponding to its specific functions. Cell morphology is mainly controlled by the combination of cytoskeletal proteins and dynamicity of plasma membrane protrusion, curvature and invagination.

Cilia, cytonemes and microvilli are representative protrusions (1). Cilia contain microtubules and act as antennae for physical stimuli or extracellular signal molecules. Cytonemes, which comprise actin, are presumed to transport the signal molecules distant from the cell body. Microvilli, which are often formed at the luminal membrane in the intestine, are also membrane protrusions rich in cholesterol (2), and are formed to widen the cell surface and to efficiently incorporate extracellular materials into the body. In general, these protrusions transduce essential information into the cells in order to decide the cell response to these stimuli. Therefore, the mechanisms for cell shape regulation is one of the central questions of cell biology.

In the vertebrate retina, the photoreceptor cell has a long cell shape, and is divided into different functional compartments. Among these compartments, the discs, which are responsible for initial light perception, are continuously formed in the outer segment. Disc formation commences with the curvature of the membrane at the adjacent region of the connecting cilium, which is then separated from the cell membrane to form microvesicles in the photoreceptor cell (3).

Prominin-1 (CD133, Prom1) encodes a pentaspan transmembrane glycoprotein, highly expressed in the retina, kidney, and testis (4). Prom1 is localised at the connecting cilium and in the outer segment (5), and is recognised as a crucial gene for the homeostasis of photoreceptor cells (5). The loss of Prom1 function leads to photoreceptor degeneration (6-8). In pedigrees with mutations in the Prom1 gene, individuals carrying the homologous mutation suffer from inherited macular dystrophies termed as Stargardt’s disease and retinitis pigmentosa (RP); the symptoms begin in childhood, followed by gradual vision loss (7-9). In our previous study, we employed Prom1 gene deficient (*Prom1*KO) mice to demonstrate that photoreceptor degeneration occurs in response to light stimulation (10). In *Prom1*KO mice, photoreceptor development and retinal structure at the perinatal stages are normal, but the membrane structure of the photoreceptor cells starts deforming once the eyes open (10). Nevertheless, as detailed molecular characterization of Prom1 is lacking, the underlying mechanisms for the initiation of retinal degeneration remain unclear.

With respect to the signalling pathway associated with Prom1, it has been demonstrated that two carboxyl cytoplasmic tyrosines of Prom1 protein are phosphorylated by the oncogenic protein kinases Fyn and Src (11). Moreover, the PI3K (phosphoinositide 3-kinase) mediated signalling pathway acts downstream of Prom1 in the glioma stem cells (12). Nevertheless, whether this activation module and the signalling pathway active in different context or the cells is elusive. Importantly, in the photoreceptor cells, PI(3) P, the product of PI3K, is predominantly localised at the inner segment, whereas Prom1 is mainly localised at the outer segment (5,13). The deletion of p85α, a subunit of PI3K, does not lead to a severe retinal degeneration (14). This suggests that the signalling pathway directly triggered by Prom1 in the photoreceptor cells is distinct from the one mediated by PI3K.

In this study, we attempted the molecular characterisation of the Prom1 protein, and identified the signalling pathway triggered by Prom1 in the retina. We found that cell morphology was considerably altered by the overexpression of Prom1 in the retinal pigmented epithelium derived cell line; numerous and long membrane protrusions, enriched in cholesterol, were formed. By using this as the evaluation criterium, we identified the essential amino acids and the downstream signalling pathway to trigger this morphological change. Importantly, chloride efflux is closely associated with the formation of the membrane protrusion. We discuss the involvement of Prom1 in membrane morphogenesis through the activity of chloride conductance.

## RESULTS AND DISCUSSION

### Membrane protrusions by Prom1 are formed independently from that of tubulin or actin polymerisation

In order to characterise the Prom1 protein, we performed a forced expression analysis of Prom1 tagged with YFP in hTERT-RPE1 (RPE1) cells. At 24 h post-transfection, we observed more than 50 membrane protrusions per cell, each with a length of more than 20 μm, on the cell surface, which were missing in the control YFP-transfected cells (Fig. 1A-C). Moreover, the overexpressed Prom1 protein was localised to these aberrantly formed protrusions (Fig. 1A).

**Fig. 1.**
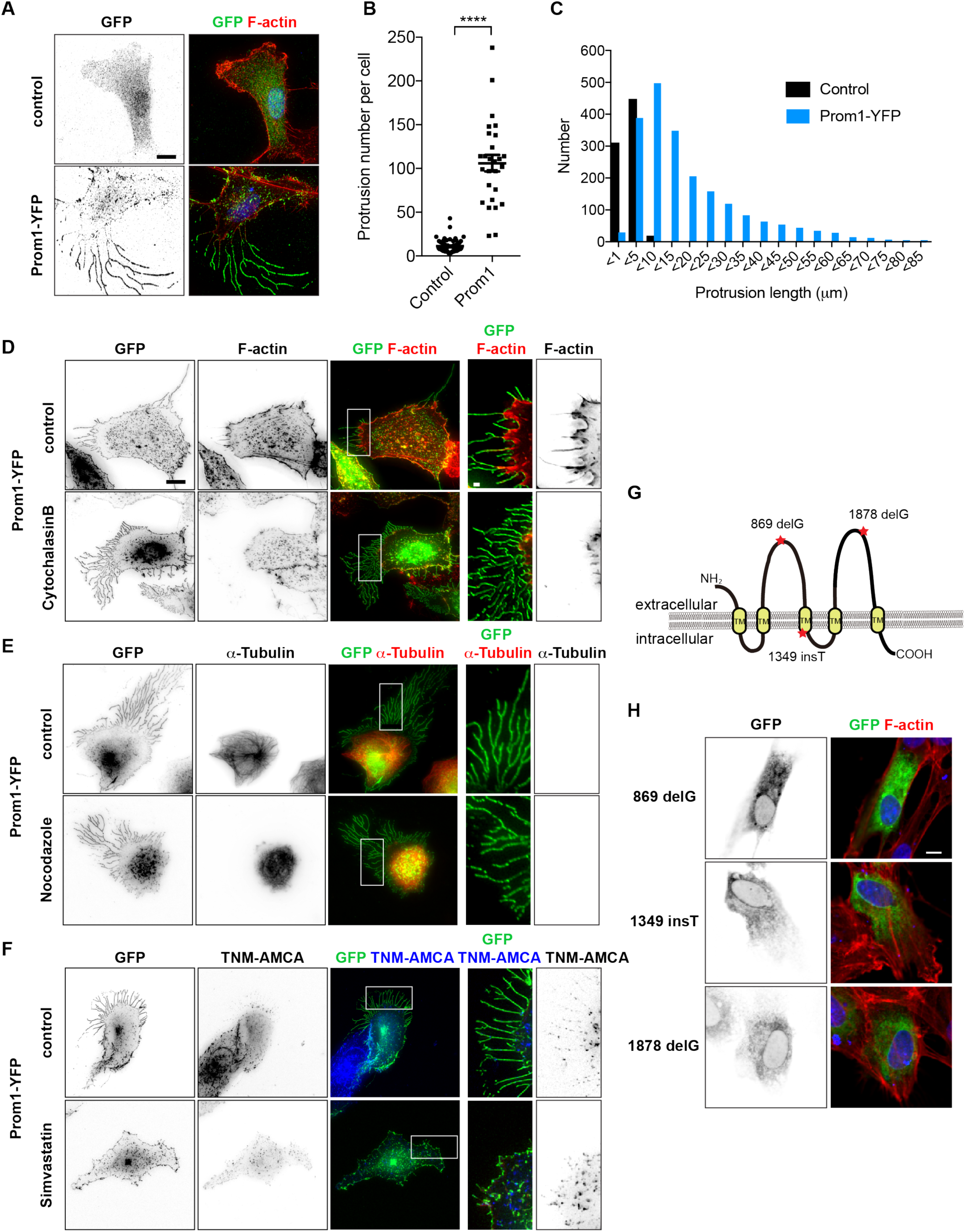
Prom1 induces cell membrane protrusions enriched in cholesterol. (A) Control *YFP* or *Prom1-YFP* were transfected into the RPE1 cells and cells were harvested to be stained with GFP antibody or phalloidin. In the high-contrasted image, yellow arrowheads indicate the tips of actin. (B,C) Quantitative data for (A). The numbers (B) and lengths (C) of the protrusions were counted and measured, respectively. (D) Cells were treated with DMSO (control), 10 μM of cytochalasin B (D), 20 μM of nocodazole (E), 1 μM of simvastatin (F) for 6 h and the expression plasmid conveying *Prom1-YFP* was transfected. At 24 h after the transfection, cells were analysed by staining with GFP (D-F) and phalloidin (C), α-tubulin (B) antibodies or TNM-AMCA (F). Enlarged images corresponding to the white squares are shown in the right panels. (G) A schematic representation of Prom1 mutations. The deletion at the 869th guanine nucleotide (869 delG), the insertion at the 1349th thymine (1349 insT) and the deletion at the 1878th guanine nucleotide (1878 delG) lead to the precocious stop codon immediately downstream of the mutation. Nucleotide count is enumerated from the start codon ATG. (H) These mutant constructs were transfected into the cells and analysed with GFP antibody and the phalloidin staining. Scale bar, 10 μm (A, D, E, F, H), 1 μm (D, E, F; two right panels).

As the protrusions often comprised actin (for cytoneme) and microtubules (for cilia) (1), we assessed whether the protrusion formation is dependent on these cytoskeletal proteins, and treated the cells with cytochalasin B and nocodazole in order to block actin polymerisation and microtubule formation, respectively. Neither of these treatments perturbed protrusion formation upon the transfection of Prom1-YFP, despite actin polymerisation (Fig. 1D) and microtubule formation (Fig. 1E) being considerably disturbed. These findings revealed that the protrusions formed by Prom1 are independent of these major cytoskeletal components with respect to both the structure and the trigger of formation.

Previous studies have reported that Prom1 is a cholesterol-binding protein (15-17). Therefore, we investigated whether cholesterol is an essential component for protrusion, and treated the cells with the cholesterol-synthesising inhibitor Simvastatin (18). The inhibitory effect was confirmed using a fluorescein sterol probe TMN-AMCA (19), and protrusion formation was completely abolished (Fig. 1F). This suggests that the cholesterol accumulation is required for protrusion formation induced by Prom1.

Various mutations have been found in the RP patients in the Prom1 gene, resulting the production of the truncated Prom1 polypeptides (Fig. 1G) (5,6,8). We therefore asked if these mutant forms of Prom1 have correlations with the protrusion formation, and overexpressed them in the cells. As the result, we found that neither of them did form the membrane protrusions (Fig. 1H), suggesting that protrusion formation and photoreceptor deformation are associated with each other.

Whilst Prom1 is known to be localised at the tips of cilia (20), the protrusions formed by Prom1 was not related to cilia (Fig. 1E). Moreover, they are enriched in cholesterol and do not require main cytoskeletal proteins for the formation (16) (Fig. 1). Thus, the Prom1 activity on cell morphology is exerted via direct rearrangement of the membrane components, without affecting on microtubule or actin.

### The five amino acids located at the carboxyl terminus are responsible for protrusion formation

Next we asked the amino acids responsible for membrane protrusion formation. Since most Prom1 mutations in the RP patients result in the production of the polypeptide lacking with its carboxyl-terminal region (Fig. 1G), we constructed a series of Prom1 truncation mutants in speculation that the responsible amino acids would reside in the carboxyl terminus (Fig. 2A).

**Fig. 2.**
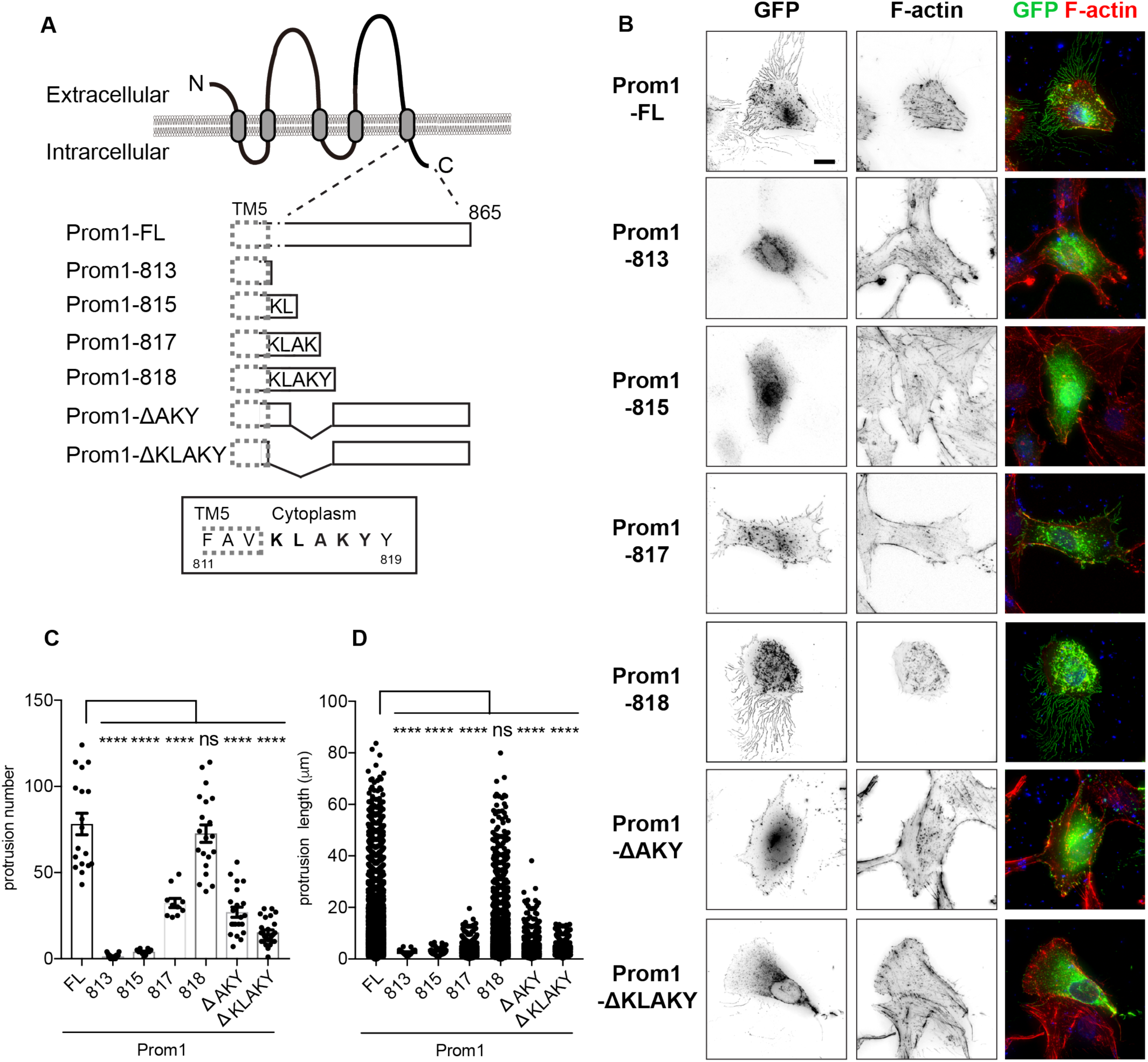
The five amino acids in the carboxyl terminal region are essential for the formation of the cell membrane protrusions. (A) A schematic representation of the Prom1 protein and its deletion mutants. (B) Representative images of the cells transfected with each deletion mutant. The expression plasmids conveying Prom1-FL (Full-length of Prom1), Prom1-813 (as indicated in (A)), Prom1-815, Prom1-817, Prom1-818, Prom1-ΔAKY or Prom1-ΔKLAKY were transfected into the cells, and the cells were analysed by staining with GFP antibody and with phalloidin at 24 hpt. Scale bar, 10 μm. (C,D) Quantitative data for (B). The numbers (C) and lengths (D) of the protrusions were counted and measured, respectively. More than 10 cells were counted and measured.

The membrane protrusion was missing in the overexpression of the Prom1 deletion mutant whose translation ends at the 813th amino acid (Fig. 2A-D). Nevertheless, when the deletion mutant that contains the five amino acids KLAKY (Lysine-Leucine-Alanine-Lysine-Tyrosine; Prom1-818) was transfected into the cells, the number and the length of the protrusion were essentially the same as those formed upon the full-length of Prom1 transfection (Fig. 2A-D), whereas the constructs that comprised a part of the KLAKY residues (Prom1-815 and 817; Fig. 2A) led to the formation of incomplete protrusions. Conversely, the construct containing the full-coding regions except for the AKY amino acid stretch form had a significantly reduced, and the construct without KLAKY had no activity to form protrusions (Fig. 2A-D). Together this finding suggests that these five amino acids are responsible for protrusion formation. We further evaluated whether the last tyrosine (Y818) requires phosphorylation for the complete activity of protrusion formation, and transfected a construct in which the tyrosine was replaced with phenylalanine (Y818F). However, protrusion formation was comparable with that with Prom1-FL (Fig. S1A). Thus, phosphorylation at this site is unlikely to be necessary for protrusion formation.

These analyses suggest that the five amino acids located immediately downstream of the fifth transmembrane domain are essential for the membrane protrusion formation.

### Rho/ROCK signalling is required for the formation of protrusions by Prom1

We next explored the essential factors that mediate the protrusion formation by Prom1. As PI3K signalling pathway (12) and the tyrosine kinases Src and Fyn (11) are essential downstream components, we observed the cell protrusion formed upon the Prom1 transfection in the cells pre-treated with LY294002 or CGP77675, pan-PI3K and Src inhibitors, respectively. However, no effect of these chemical treatments on protrusion formation was observed (Fig. S1B,C). Moreover, the substitution mutant Y828F, which abolishes the essential phosphorylation for the Src signalling activation (12), was as active as Prom1-FL regarding the protrusion formation (Fig. S1D). This observation suggests that Prom1 has distinct downstream branches, and the membrane protrusions formed by Prom1 are induced via differing signalling mediator(s) from those previously reported.

We therefore screened the downstream signalling of Prom1 by evaluating protrusion formation following treatment with signal inhibitors. We specifically emphasized the inhibitors of the small GTPases, including Rho, Rac and Cdc42, as these GTPases are often involved in membrane protrusion formation (21).

While EHT1864 and ZCL279, selective inhibitors for Rac1 and Cdc42, respectively, did not have an effect on protrusion formation by Prom1 (Fig. 3A-C), we observed that the ROCK inhibitor Y-27632 substantially reduced the number and the length of the protrusions (Fig. 3A–C). As the ROCK inhibitor affects both Rho and Rac, we further attempted to identify the molecule, and used C3, a membrane-permeable recombinant protein that specifically blocks the Rho signal. C3 had a similar effect to that of Y-27632 (Fig. 3A–C); protrusion formation was profoundly blocked. Consistently, the co-transfection of the dominant-negative RhoA in combination with Prom1 blocked protrusion formation (Fig. 3D). These findings suggest that the Rho-associated signalling pathway is essential in the membrane protrusion formation by Prom1.

**Fig. 3.**
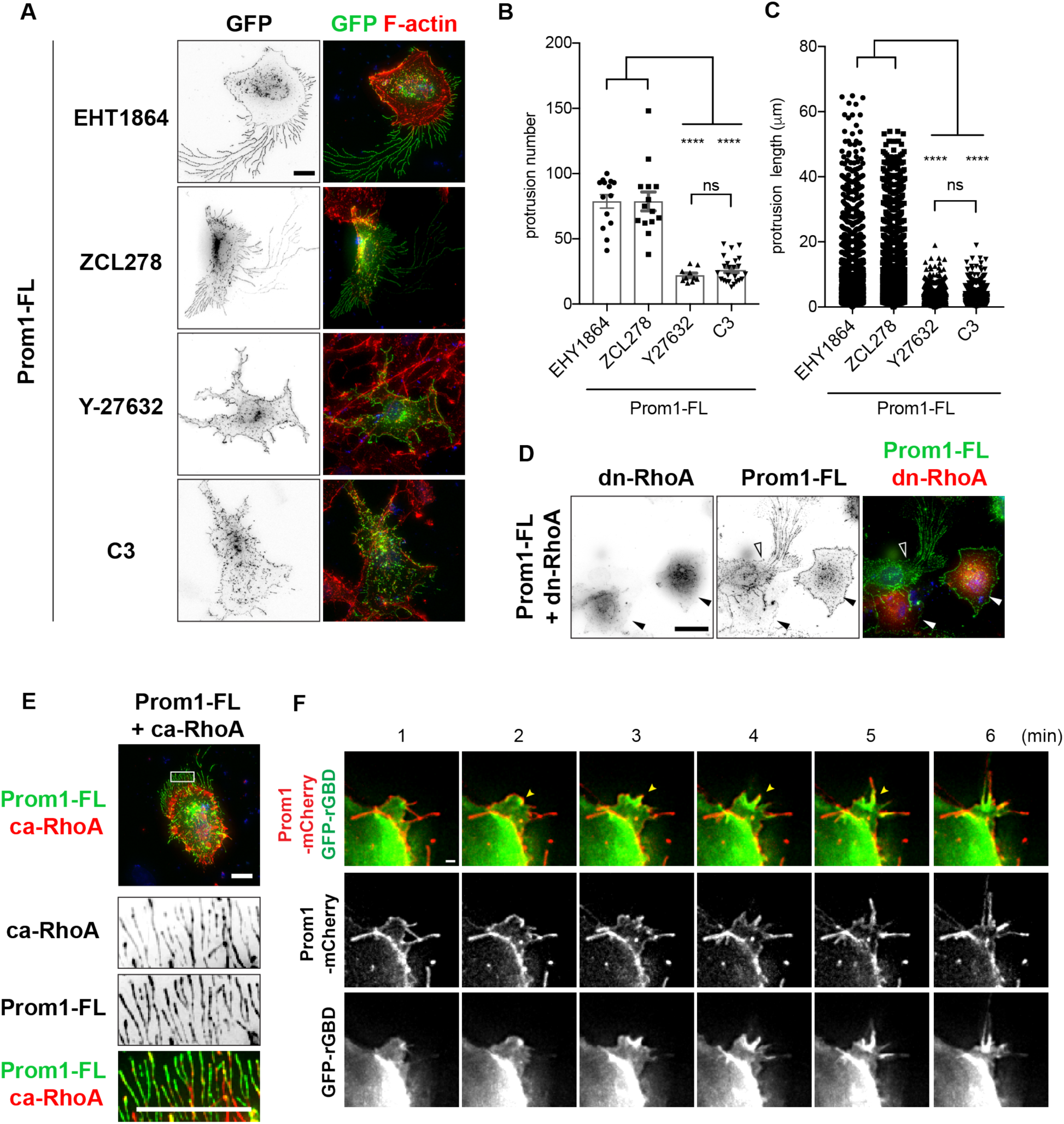
RhoA is essential for protrusion formation induced by Prom1. (A) The inhibitors targeting the small GTPases, as indicated, were treated before *Prom1-YFP* was transfected. Scale bar, 10 μm. (B,C) Quantitative data for (A). The numbers (B) and lengths (C) of the protrusions were counted and measured, respectively. (D) The plasmid conveying *myc-tagged dominant-negative version of RhoA* (*dn-RhoA*) was co-transfected with *Prom1-FL*. Staining was performed with GFP (for Prom1), myc (for dn-RhoA) antibodies and phalloidin. (E) The plasmid conveying *myc-tagged constitutively-active version of RhoA* (*ca-RhoA*) was co-transfected with *Prom1-FL*. Staining was performed with GFP (for Prom1), myc (for dn-RhoA) antibodies and phalloidin. Enlarged images corresponding to the white squares are shown in the bottom three panels. (F) The plasmids conveying *GFP-rGBD* and *Prom1-mCherry* were co-transfected and time-lapse imaging was performed for 6 minutes at 24 hpt. Yellow arrowheads show the protruding membrane. Scale bar, Scale bar, 10 μm (A, E), 20 μm (D), 1 μm (F).

Conversely, as revealed by the transfection of the constitutively-active form of RhoA, the active Rho was co-trafficked into the protrusions formed by Prom1 (Fig. 3E). Eventually, we highlighted the initial moment when the protrusion was formed. We co-transfected GFP-rGBD (22), which visualises the activated Rho, together with Prom1-mCherry into the cells, and evaluated the individual proteins via time-lapse analysis. We found that Rho was activated at the plasma membrane, and protrusion formation was initiated at the membranous point where the active Rho and Prom1 were encountered (Fig. 3F and movie S1). Collectively, these findings suggest that the membrane protrusion formed by Prom1 is mediated by the small GTPase RhoA.

Protrusion formation by Prom1 requires co-localization with active Rho (Fig. 3). Nevertheless, according to the immunoprecipitation analysis, Prom1 does not bind to or activate Rho (Fig. S2). This suggests that Rho is activated by another triggering factor(s), including specific RhoGEFs (Rho family specific GDP-GTP guanine exchanging factors), and interacts with Prom1 weakly or transiently.

### Prom1 drives the chloride ion efflux upon the intracellular calcium ion uptake

The high-dimensional structure-based homology search algorithm HHPred (23) predicted that Prom1 is highly homologous with the membrane proteins TTYH1/2 (Fig. S3A). (24). The overexpression of TTYH2 in the RPE cells induced membrane protrusions in a manner similar to Prom1 (Fig. S3B), suggesting that Prom1 and TTYH2 have functional similarities. As the TTYH-type receptors are known to act on the calcium-activated chloride currents (25), we hypothesised that Prom1 has a similar function.

To address this question, we used mouse embryonic fibroblast (MEF) cells extracted from wild-type or *Prom1* gene-deficient (*Prom1* knockout; *Prom1*KO) embryos (10,26), and measured the temporal change of the intracellular chloride ion level upon calcium uptake by using the chloride-sensitive fluorescent indicator MQAE (N-(ethoxycarbonylmethyl)-6-methoxyquinolinium bromide). As MQAE is quenched by chloride ion, the fluorescein intensity is reciprocal to the intracellular chloride ion concentration. Once the intracellular calcium uptake was provoked by the calcium ionophore A23187, significant chloride efflux was observed in the wild-type cells within a several minutes (8 min; Fig. 4A,B and movie S2A). In contrast, the extent of the efflux was reduced approximately by 50% in the *Prom1*KO cells (8 min; Fig. 4A,B and movie S2B), suggesting that the chloride ion was accumulated in the cells. A similar result was obtained in another analysis in which the cell mass was measured (Fig. S4A). Importantly the extent of the calcium uptake upon the A23187 treatment was comparable (Fig. S4B), suggesting that the perturbation of chloride ion efflux was not the secondary effect caused due to the change in calcium influx. Collectively, these observations suggest that Prom1 modulates the dynamic intracellular chloride current upon the calcium uptake.

**Fig. 4.**
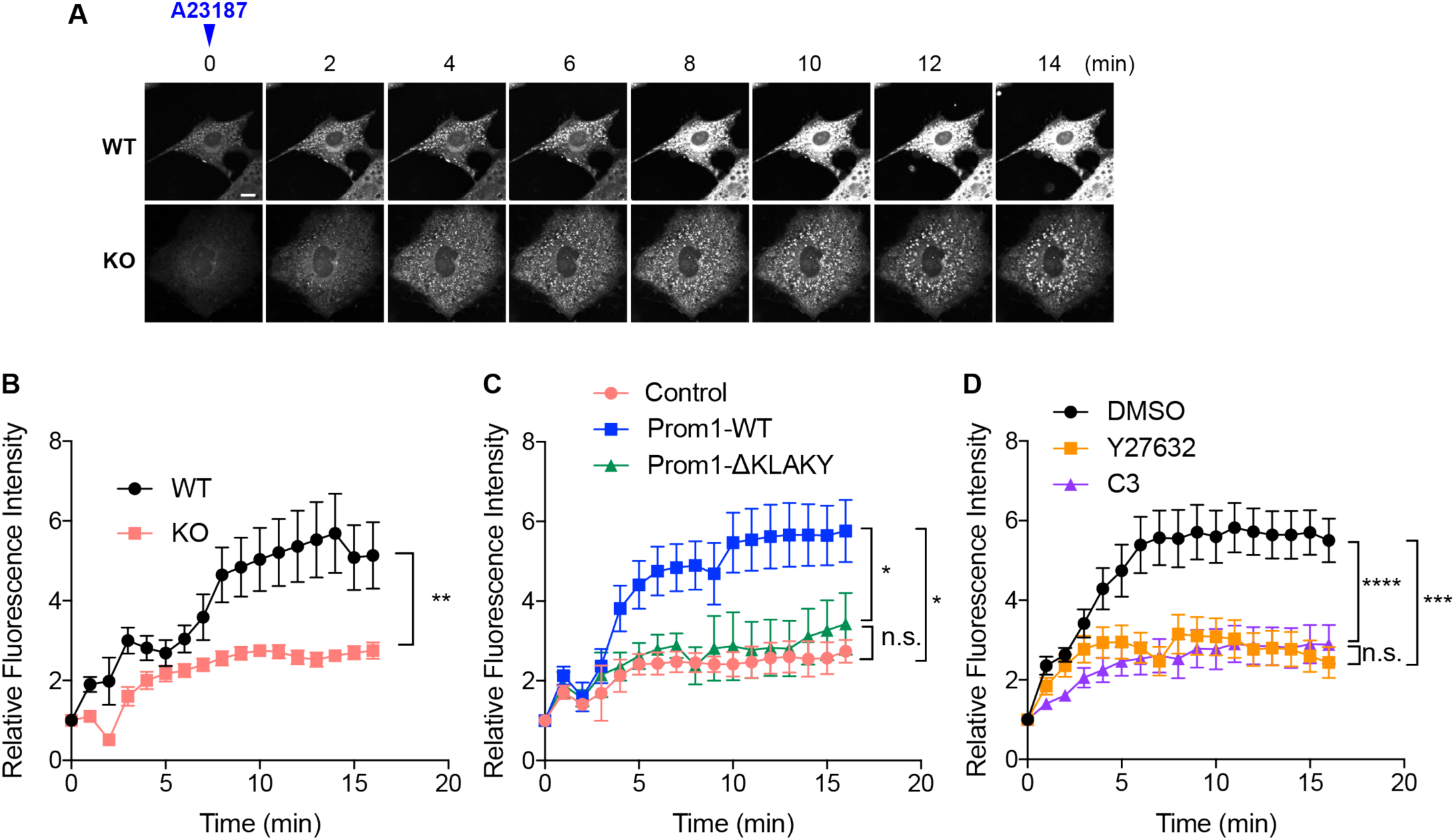
Prom1 modulates the chloride conductance upon intracellular calcium uptake. (A) The temporal change in fluorescein intensities of MQAE was measured. The wild-type and *Prom1*KO MEF cells were incubated with low-chloride Kreb’s medium (see materials and methods) and the intracellular calcium uptake was provoked by adding 5 μM of the calcium ionophore A23187 onto the medium (time 0). The temporal change of the fluorescein intensity was imaged at 1 min intervals up to 15 min after the ionophore treatment under the confocal microscope. Representative images are presented. Scale bar, 10 μm. (B) Quantitative data for (A). Eight cells were selected from each of wild-type and *Prom1*KO cells, and the fluorescein intensities at each time point were quantified. Data are represented as the mean values ± s.e.m. (C) The expression plasmids conveying *Prom1-FL* or *Prom1-ΔKLAKY* were transfected into the *Prom1*KO MEF cells, and cells were incubated in the presence of MQAE. The transfected cells were identified by YFP expression, and the fluorescein intensities from each transfection were traced. (D) The chloride efflux is perturbed upon the treatment with Rho inhibitors Y27632 and C3. Wild-type MEF cells were treated with 20 μM of Y-27632 or with 0.5 μg/ml of C3 for 2 hours at the same time of the MQAE treatment and were subjected to the fluorescein measurement as in (B) and (C).

Furthermore, we investigated whether this efflux perturbation in the *Prom1*KO cells was rescued upon the transfection of the wild-type Prom1 and and Prom1-ΔKLAKY. When we transfected Prom1-FL in the *Prom1*KO cells, the chloride efflux was found to be restored to the same level as in the wild-type MEF cells (7 cells; Fig. 4C, Fig. S5A and movie S3). However, this outflow failed to occur upon the transfection of Prom1-ΔKLAKY, suggesting that the amino acids stretch KLAKY (Fig. 2A) was essential for the regulation of the chloride efflux. Moreover, the wild-type MEF cells pre-treated with Rho/ROCK inhibitors Y-27632 or C3 perturbed the chloride efflux upon the calcium uptake (Fig. 4D, Fig. S5B and movie. S4). Collectively, these findings suggest that the function of Prom1 as the membrane morphology modulator and the chloride ion current regulator are closely associated with each other.

In this study, we demonstrated that Prom1 induces the formation of membrane protrusions enriched with cholesterol, and this activity is dependent on the carboxyl-termnal domain of the protein. We also demonstrated that Prom1 is structurally similar to TTYHs, proteins involved in the calcium-activated chloride currents (25,27), and it is involved in the chloride current activated by calcium uptake (Fig. 4). While the physiological significance of TTYHs in the retinal homeostasis remains unclear, ionic current in the photoreceptor cells is apparently crucial for their functions (28). In physiological level, one major protein that uptakes the intracellular calcium ion is rhodopsin. Rhodopsin is a GPCR (G-protein coupled receptor) that converts light stimuli to the cGMP activation followed by the intracellular calcium uptake (29). In our immunoprecipitation analysis, rhodopsin interacted with Prom1 (Fig. S6), suggesting that these two proteins act in conjugation with each other. As rhodopsin is activated by light stimuli, it can induce Prom1 activity, and protrusion formation and chloride current may occur. In *Prom1*KO mice, the outer segment of the photoreceptor cells is not appropriately formed (9,10). Moreover, as it has been reported that the newly formed discs are enriched in cholesterol (30). Therefore it is reasonable to speculate from our data that Prom1 controls the evagination of the newly formed disc by interacting with cholesterol.

Future analyses, including single photoreceptor recordings of the temporal change of chloride ion and the membrane evagination in wild-type and *Prom1*-mutant cells, will identify the initial step of the photoreceptor degeneration and will provide new insight in developing novel therapeutic methods for intractable hereditary retinopathies.

## Materials and Methods

### Ethical statement on animal experiments

All animal experiments were carried out with the approval of the animal welfare and ethical review panel of Nara Institute of Science and Technology (approval numbers: 1533 and 1810 for animal research, and 311 for genetic modification) and Institutional Animal Care and Use Committee of RIKEN Kobe branch. *Prom1*KO mice established previously (26) (CDB0623K: http://www2.clst.riken.jp/arg/methods.html) were reared as a hybrid genetic background of C57BL/6 and CBA/NSlc (10).

### Cell culture, transfection and Rho activation assay

hTERT-RPE1 (ATCC CRL-4000) was cultured in high-glucose Dulbecco’s Modified Eagle Medium (DMEM; Wako, Japan) containing 10% FBS (Gibco) supplemented with non-essential amino acids, glutamine and penicillin/streptomycin (Wako, Japan).

While multiple isoforms have been reported for the Prom1 transcripts (4,31), we employed the isoform encoding 865 amino acid. The Prom1 constructs were carboxyl-terminally fused with YFP or mCherry as indicated. DN-Rho and ca-Rho were constructed as described previously (32).

The plasmids were transfected with Lipofectamine-2000 (Invitrogen). Rho activation assay was performed by using the Rho activation assay kit (Millipore). Immunoprecipition was performed with the magnetic beads conjugated with myc antibody.

Antibodies used in this study were; GFP (rabbit; MBL; #598), myc (mouse; CST; #2276S), HA (mouse; SIGMA; #H9658). Chemicals were; cytochalasin B (Wako, Japan; #030-17551), nocodazole (SIGMA; #M1404), Simvastatin (Cayman chemical; #10010344), EHT1864 (Cayman chemical; #17258), ZCL278 (TOCRIS; #4794), Y-27632 (Wako, Japan; #251-00511), C3 (Cytoskeleton, Inc; #CT04). LY294002 (Wako, Japan; #129-04861), CGP77675 (Cayman Chemical; #21089).

### Immunofluorescence microscopy and protrusion analysis

Immunofluorescence microscopy was performed as described previously (33). Fluorescence microscopic analyses were carried out using DeltaVision Elite Microscopy System (GE Healthcare, UK). Z-axial images were taken at 0.2 μm with a 40X objective lens. Deconvolution of images was performed using DeltaVision SoftWoRx software. Captured images were processed with Adobe Photoshop CS5. The numbers and lengths of protrusions formed on the cell membrane were measured with ImageJ software and at least 20 cells were analysed on each experiment.

### Intracellular chloride ion measurement on MEF cells

Mouse embryonic fibroblasts (MEF) were prepared from 14.5 dpc (days post-coitum) mouse embryos as described previously (34). For measuring the intracellular chloride ion level, the chloride-sensitive fluorescent indicator MQAE (Dojindo) was used and was used to treat the MEF cells according to the manufacturer’s instruction. Briefly, MEF cells were cultured in the low-chloride medium (Krebs-HEPES buffer; 20 mM HEPES-NaOH (pH 7.3), 128 mM NaCl, 2.5 mM KCl, 2.7 mM CaCl_2_, 1 mM MgSO_4_, 16 mM glucose) and the final concentration of 5 mM of MQAE was added, along with measurement of the basal chloride level. The calcium ionophore (A23187; Sigma) was then added at 5 μM and the temporal change in the chloride ion was measured using LSM 710 confocal microscope (Zeiss) or with the plate reader Tristar2 (Berthold Technologies) at 1 min intervals.

### Structure prediction, images, and data analysis

The homology search based on the secondary structure was conducted using the prediction algorithm HHPred (23). Images were observed using LSM 710 confocal microscope (Zeiss) or DeltaVision Elite (GE Healthcare) and processed by the Photoshop software (Adobe). Statistical analysis was performed by two-tail t-test using the Prism software (graphpad.com) and *p*-values (*; *p* < 0.05, **; *p* < 0.01, ***; *p* < 0.001) are indicated in each graph.

## Acknowledgements

The authors are grateful to Dr. Shinichi Nishimura for TNM-AMCA, Prof. Takashi Toda for encouragement and all laboratory members for support and valuable discussion. GFP-rGBD was distributed from Addgene, which had been deposited by Dr. William Bement (Addgene plasmid # 26732).

## Author Contributions

NS and TK conceived the project. KN initially found the phenotype induced by Prom1, and MJH predicted that Prom1 is involved in ion flux. AH performed the majority of the experiments and analysed the data with assistance from YY, KH, WK and YI. HK, SO and KK contributed to the establishment and maintenance of the *Prom1*-deficient mice. All authors joined the discussion. NS drafted and AH edited the manuscript.

