## supplementary information for "Membrane Protrusion Formation Mediated by Rho/ROCK Signalling and Modulation of Chloride Flux"

List of material included:
Supplementary Figure S1-S6
Supplementary Movies S1-S4

**Fig. S1 PI3K and src and their related amino acids in Prom1 are not involved in the cellular morphogenesis generated by Prom1.** The Prom1 mutants in which the 818th and 828th tyrosines were replaced with phenylalanine (Y818F; A, Y828F; D) were transfected. PI3K and src inhibitors do not have the inhibitory effects on the membrane protrusions generated by Prom1. 10 μM of LY294002 (a

PI3K inhibitor; B) or and 2 μM of CGP77675 (a src inhibitor; C) were treated for 6 h and Prom1-FL was transfected. In all cases, the cells were harvested 24 h after the transfection and the cell shape was analysed by the GFP antibody and phalloidin staining. Scale bar, 10 μm.

**Fig. S2 Prom1 does not activate or interact with Rho.** (A,B) Rho activation assay. The empty vector (EV) or the plasmid conveying *Prom1* were transfected into the RPE-1 cells and Rho-activation assay was performed at 24 (A), 8, or 16 hpt (B). (C) Prom1 does not interact with RhoA. Plasmids conveying *Prom1-YFP*, *Rho-myc* and the control vector (empty vector; EV) were transfected as indicated, and immunoprecipitation was performed with the magnetic beads conjugated with myc antibody and detected with the GFP antibody.

**Fig. S3 Prom1 is structurally analogous with TTYH proteins, and some of CaCC proteins have activities to induce the membrane protrusions, as Prom1 does.** (A) The outcome of the homology search by using the algorithm HHPred. (B) TTYH2 induces the membrane protrusions. The expression plasmids encoding *TTYH2* were transfected and the cell morphology was observed 24 h after the transfection. Scale bar, 10 μm.

**Fig. S4 The fluorescein measurement from a plate leader demonstrates similar results as in Fig. 4A, and the intracellular calcium uptake induced by A23187 is comparable in the wild-type and *Prom1*KO cells.** (A) The MQAE fluorescein intensities emitted from the cell mass were measured. $1 \times 10^3$ cells of wild-type or *Prom1*KO MEF cells were plated on a 96-well plate, and were treated with 5 μM of A23187. The measurement was performed at 1 min intervals for 20 min. The measurements were performed thrice for both wild-type and the *Prom1*KO cells and data are represented as the mean values ± s.e.m. (B) Intracellular calcium uptake is not affected in the *Prom1*KO cells upon the treatment with A23187. The experiment was performed as in (A), except that the cells were treated with Fluo-4, a fluorescent calcium indicator.

**Fig. S5 Typical images of temporal fluorescein changes during the culture with MQAE.** The typical images producing Fig. 4B-D are presented. Scale bar, 10 μm.

**Fig. S6 Rhodopsin and Prom1 interact with each other and are co-localised in the protrusion.**
(A) Localisation of rhodopsin in the protrusion. The expression plasmids conveying *Rhodopsin-HA* and *Prom1-YFP* were cotransfected into the RPE-1 cells and were harvested to be analysed via immunocytochemistry using HA and GFP antibodies. Scale bars, 10 μm. (B) Both Prom1 and Prom1-ΔKLAKY physically interact with rhodopsin. Prom1-FL and rhodopsin-HA were transfected as

indicated. Immunoprecipitation and was performed with the magnetic beads conjugated with HA, and the expression was detected with GFP and HA antibodies.

**Supplementary Movies**

Supplementary Movie S1 The continuous pictures related to Fig. 3F were processed to the video.

Supplementary Movie S2 The continuous MQAE images on wild-type (A) and *Prom1*KO cells (B), related to Fig. 4A were processed to the video.

Supplementary Movie S3 The continuous pictures on GFP (control GFP (A), Prom1FL (C) and Prom1ΔKLAKY (E)) and MQAE images (GFP (B), Prom1FL (D) and Prom1ΔKLAKY (F)), related to Fig. 4C were processed to the video.

Supplementary Movie S4 The continuous MQAE images with DMSO (A), Y-27632 (B) and C3 (C), related to Fig. 4D were processed to the video.
